# Exploring Efficient Linear Mixed Models to Detect Quantitative Trait Locus-by-Environment Interactions

**DOI:** 10.1101/2020.07.25.220913

**Authors:** Eiji Yamamoto, Hiroshi Matsunaga

## Abstract

Genotype-by-environment interactions (G×E) are important for understanding genotype–phenotype relationships. To date, various statistical models have been proposed to account for G×E effects, especially in genomic selection (GS) studies. Generally, GS does not focus on the detection of each quantitative trait locus (QTL), while the genome-wide association study (GWAS) was designed for QTL detection. G×E modeling methods in GS can be included as covariates in GWAS using unified linear mixed models (LMMs). However, the efficacy of G×E modeling methods in GS studies has not been evaluated for GWAS. In this study, we performed a comprehensive comparison of LMMs that integrate the G×E modeling methods to detect both QTL and QTL-by-environment interaction (Q×E) effects. Model efficacy was evaluated using simulation experiments. For the fixed effect terms representing Q×E effects, simultaneous scoring of specific and non-specific environmental effects was recommended because of the higher recall and improved genomic inflation factor value. For random effects, it was necessary to account for both G×E and genotype-by-trial (G×T) effects to control genomic inflation factor value. Thus, the recommended LMM includes fixed QTL effect terms that simultaneously score specific and non-specific environmental effects and random effects accounting for both G×E and G×T. The LMM was applied to real tomato phenotype data obtained from two different cropping seasons. We detected not only QTLs with persistent effects across the cropping seasons but also QTLs with Q×E effects. The optimal LMM identified in this study successfully detected more QTLs with Q×E effects.

## INTRODUCTION

Phenotypes are determined not only by genetic potential but also by environmental growth conditions. More specifically, the rank of phenotypic values often changes when the same genotype set is phenotyped under different environmental conditions (Cooper and DeLacy 1994). The phenotypic response to the environment is explained by reaction norms that describe the pattern of phenotypic expression of a genotype across different environments. Genotype-by-environment interactions (G×E) occur when the slopes of the reaction norms of two different genotypes are not parallel across environments (Malosetti *et al*. 2013). G×E analysis is important for a precise understanding of genotype–phenotype relationships as well as for the design of crop varieties that fit a given environment.

To date, various statistical approaches have been proposed for G×E analysis (reviewed in Malosetti *et al*. 2013). The simplest approach is analysis of variance, which compares the mean and variance of the phenotypic values of genotypes in multiple environments. However, more flexible methods were eventually developed, beginning with the <u>A</u>dditive <u>M</u>ain effects and <u>M</u>ultiplicative <u>I</u>nteraction (AMMI) model, which divides the genetic contribution for a phenotypic value into additive main effects (i.e., genetic effects not specific to the environment) and G×E effects (Gauch 1988).

Thus, G×E effects in the AMMI model are expressed as multiplicative interaction terms consisting of the products of genotypic and environmental scores. Next, principal component (PC) analysis of these scores is conducted (Gauch 1988). This approach allows graphical representation of the interactivity between genotypes and environments (Gauch 1988). In particular, a genotype will show the highest performance in the closest environment in the biplot. The <u>G</u>enotype main effects and <u>G</u>×<u>E</u> (GGE) model resembles the AMMI model, but is focused on the total genetic effect in each environment (Yan *et al*. 2000). In the GGE model, both additive main effects and environment-specific genetic effects are scored together; thus, genotypes in the GGE biplot represent the overall genetic effect, while the AMMI biplot represents only the G×E effect (Yan *et al*. 2000). The AMMI and the GGE models provided an important perspective on G×E analysis, and facilitated discussion about whether genetic effects for a phenotypic value within a given environment should be separated into specific and non-specific environmental effects.

In the AMMI and GGE models, genotypic differences are treated as categorical variables (Gauch 1988; Yan *et al*. 2000). However, the availability of molecular markers has allowed the quantitative description of genotypic differences using a <u>g</u>enomic <u>r</u>elationship <u>m</u>atrix (GRM) (Endelman and Jannink 2012). GRM applications in G×E analysis have advanced considerably in genomic selection (GS) studies (reviewed in Crossa *et al*. 2017). In GS, a training population that have been phenotyped and genotyped is used to construct a model that predicts the genetic potential of unphenotyped individuals (Meuwissen *et al*. 2001). Most GS models that account for G×E effects are constructed as linear mixed models (LMMs) (Crossa *et al*. 2017). LMMs treat genetic effects as random effects using a variance–covariance matrix designed using GRMs. The most straightforward method is to consider G×E random effects as a lack of genetic correlation between environments (Lopez-Cruz *et al*. 2015). If information on the similarity among environments is available as a variance–covariance matrix, then G×E random effects can be designed as the product of a GRM and the variance–covariance matrix (Cuevas *et al*. 2017). If there are multiple options to model random effects, then it may be preferable to integrate all random effect terms into a single model and estimate their relative contributions to phenotypic values (Jarquín *et al*. 2014). In all of these methods, random effects with G×E have common variance across environments (Lopez-Cruz *et al*. 2015; Cuevas *et al*. 2017). Sousa *et al*. (2017) extended the models to allow different variances across environments, which resulted in increased GS prediction accuracy in some experiments.

Generally, GS models do not test the significance of each marker effect because their objective is phenotype prediction and not the detection of quantitative trait loci (QTLs) (Meuwissen *et al*. 2001; Crossa *et al*. 2017). For QTL detection, genetic mapping approaches such as the genome-wide association study (GWAS) are necessary (Cortes *et al*. 2020). Currently, the most frequently used GWAS method is based on unified LMMs (Yu *et al*. 2006). In the unified LMMs, total genetic effects are divided into a fixed effect for each marker genotype and random effects that are modeled for all marker genotypes (Yu *et al*. 2006). Then, the statistical significance of the fixed effect is analyzed to estimate the QTL in linkage disequilibrium (LD) with the marker.

Because the methods used to model G×E in GS studies were designed for random effects (Jarquín *et al*. 2014; Lopez-Cruz *et al*. 2015; Cuevas *et al*. 2017; Sousa *et al*. 2017), they can be applied to random effects in unified LMMs for GWAS. Although several studies have detected QTL-by-environment interaction (Q×E) effects using LMMs (Boer *et al*. 2007; Mathews *et al*. 2008), the efficacy of G×E random effects methods in GS studies for Q×E analysis has not been surveyed, because this is a recent application. Therefore, the advantages and disadvantages of the various methods developed to model G×E random effects in Q×E analysis are poorly understood. In the present study, we performed a comprehensive comparison of the methods used to model G×E random effects in GWAS. The objective of this study was to identify the most effective LMMs for detecting QTLs with Q×E effects. For fixed QTL effects, we compared two methods that were designed based on ideas represented in the AMMI (Gauch 1988) and the GGE (Yan *et al*. 2000) models. The first method divides the total genetic effects of a QTL in LD with a marker into two effects: the additive main effect, which is not specific to the environment, and the Q×E effect. The second method simultaneously scores the additive main effect and the Q×E effect. All combinations of the modeling methods for fixed effect terms and random effect terms were compared using simulation experiments. Finally, LMMs selected based on the simulation experiments were applied to GWAS using tomato (*Solanum lycopersicum* L.) phenotype data.

## MATERIALS AND METHODS

### Assumptions

To clarify the aim of the present study and its differences from similar recent studies (e.g., Moore *et al*. 2019; Dahl *et al*. 2020), we describe our assumptions as follows:

#### Experimental design

- The environment in this study can be geographic location, weather condition, and other artificial experimental conditions. In the real phenotype data used in this study, cropping season was the environmental differences (Figure 1).
- Differences among environments and/or trials are represented as categorical variables.
- Phenotypic values of a genotype in an environment were obtained for several trials (Figure 1).

**Figure 1.**
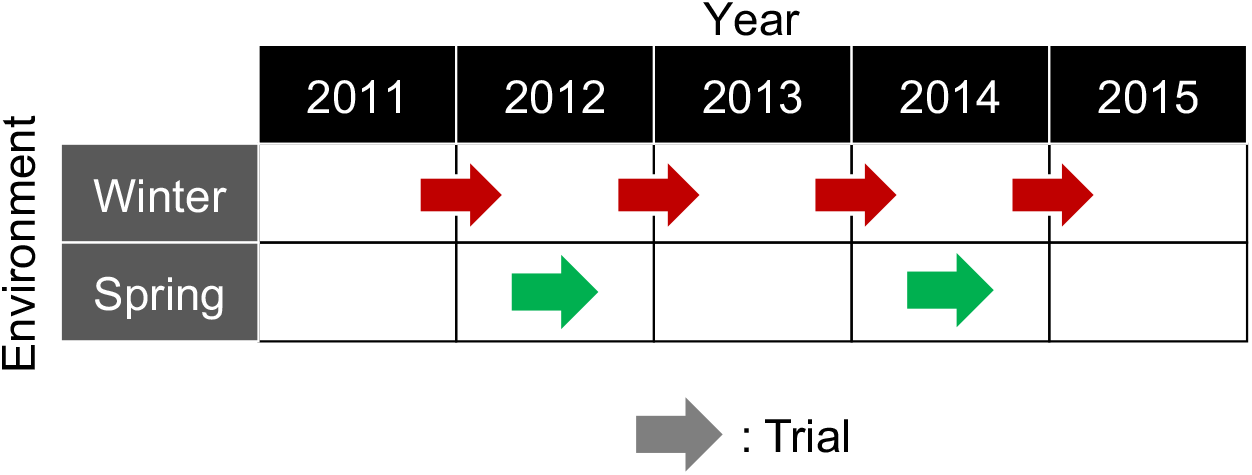
Environments and trials used in this study. Cropping season (i.e., winter or spring harvest cropping seasons) was the environmental variable in the tomato phenotypic dataset. Each trial represents a period of plant growth and phenotype evaluation.

#### Genetic effects

- The target traits are controlled by several major QTLs and numerous minor QTLs, and the major QTLs were the detection target.
- Genetic effects in the same environment are affected by genotype-by-trial (G×T) effects.

### LMMs

We modeled the effects of Q×E and G×E using unified LMMs, in which the total genetic effects were divided into fixed and random effect terms:

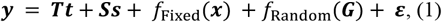

where ***y*** and ***ε*** indicate *n* × 1 vectors for phenotypic values and residuals, respectively; ***T*** is an *n* × *TR* design matrix that assigns phenotypic values to trials, where *TR* is the number of trials; ***t*** is a *TR* × 1 vector of population-wide mean for each trial; ***S*** is a *n* × *c* matrix whose column elements are the eigenvectors from principal component analysis of genotype data from all markers. ***s*** indicates a *c* × 1 vector of fixed effects for ***S. S*** and ***s*** are used to decrease the rate of false signals generated by the population structure (Price *et al*. 2006; Yang *et al*. 2011; Li *et al*. 2014). In our analyses of simulations and real data in this study, ***S*** consisted of the first and second eigenvectors, which explained 30.9% and 7.8% of the total genetic variation, respectively (Yamamoto *et al*. 2016). *f*_Fixed_(***x***) represents the fixed effect terms for a major QTL in LD with a single-nucleotide polymorphism (SNP) marker. *f*_Random_(***G***) represents the random effect terms for the background genetic effects. ***x*** is an *n* × 1 vector of SNP genotype values coded as {0, 1, 2} = {aa, Aa, AA}; ***G*** is an *m* × *m* GRM containing *m* genotypes. In the GRM, the genetic relationship between individuals *j* and *k*(*G_jk_*) is defined as:

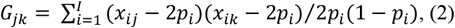

where *x_ij_* is the coded genotype value for the *i*-th SNP of the *j*-th individual, *p_i_* is the minor allele frequency for the *i*-th SNP, and *I* is the total number of markers. The GRM was calculated using the *A. mat* function in the R package “rrBLUP” (Endelman and Jannink 2012). Because the effects of major QTL and the Q×E effects were included in *f*_Fixed_(***x***), the proportion of variance explained by *f*_Fixed_(***x***) was subjected to a statistical test, whereas *f*_Random_(***G***) and the other terms were covariates. Detailed descriptions of *f*_Fixed_(***x***) and *f*_Random_(***G***) are provided in the sections “Modeling fixed effects” and “Modeling random effects”, respectively. In a standard unified LMM (Yu *et al*. 2006), Eq. 1 is described as follows:

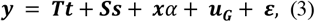

where *α* models the additive main effect of a QTL in LD with the SNP. We refer to the fixed QTL effect term in Eq. 3 (i.e., ***x**α*) as the additive main effect term (Table 1). ***u_G_*** models the random effects as follows:

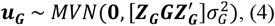

where *MVN* is the multivariate normal distribution; ***Z_G_*** represents an *n* × *m* incidence matrix for the phenotype and random effects; and 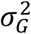 is the variance for ***u_G_***.

**Table 1.** Terms used for the linear mixed models examined in this study.

| Terms | Equation in<br>main text | Definition |
| --- | --- | --- |
| Fixed effects |  |  |
| Additive main | Eq. 3 | No Q×E effect assumed. |
| AMMI-type Q×E | Eq. 5 | QTL effect divided into additive main and Q×E effects. |
| GGE-type Q×E | Eq. 6 | Additive main and Q×E effects scored together. |
| Random effects |  |  |
| $\mathbf{u}_G$ | Eq. 7 | No G×E random effects assumed. |
| $\mathbf{u}_{GE}$ | Eq. 8 | G×E random effects not correlated between environments but have common variance. |
| $\mathbf{u}_{GT}$ | Eq. 10 | G×T random effects not correlated between environments but have common variance. |
| $\sum_{tr}^{TR} \mathbf{u}_{tr}$ | Eq. 13 | G×T effects not correlated between trials and have different variances. |

AMMI, Additive Main effects and Multiplicative Interaction model; G×E, genotype-by-environment interaction; GGE, Genotype main effects and G×E model; G×T, genotype-by-trial interaction; Q×E, QTL-by-environment interaction; QTL, quantitative trait locus.

#### Modeling fixed effects

To model Q×E as fixed effects in the LMMs, we used two formulae. The first is as follows:

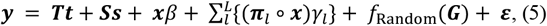

where *β* is the QTL effect not specific to the environment, and *γ_l_* is the QTL effect specific to the *l*-th environment. ***π**_l_* is an *n* × 1 vector containing indicator variables that determine whether the phenotypic value is obtained from *l*-th environment {1} or not {0}. The symbol ∘ indicates the Hadamard product for the left and right vectors or matrices. In this equation, ***x**β* is analogous to the additive main genetic effect in the AMMI model (Gauch 1988). Therefore, we refer to the fixed QTL effect terms for an SNP in Eq. 5 (i.e., 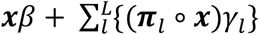) as AMMI-type Q×E effect terms (Table 1). The second formula used to model Q×E is as follows:

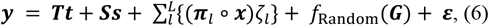

where *ζ_l_* is the QTL effect in the *l*-th environment. Unlike Eq. 5, Eq. 6 does not include the QTL effect not specific to the environment (i.e., ***x**β*). Therefore, the Q×E effect terms are analogous to the GGE model (Yan *et al*. 2000). We refer to the Q×E effect terms in Eq. 6 (i.e., 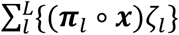) as GGE-type Q×E effect terms (Table 1).

#### Modeling random effects

Appropriate statistical modeling for random effects has been the key to efficient GWAS using LMMs (Yu *et al*. 2006). In a standard GWAS, random effects are modeled as in Eq. 4. Therefore,

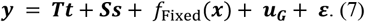

Notably, ***u_G_*** does not account for the G×E effects (Yu *et al*. 2006). Therefore, we extended Eq. 7 as follows:

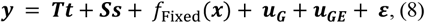

where ***u_GE_*** models the G×E effects as follows:

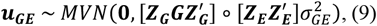

where ***Z_E_*** is the incidence matrix for the phenotypic values and environmental differences (e.g., cropping season in Figure 1). Thus, Eq. 9 allows for different random effects among environments. The third random effect model is as follows:

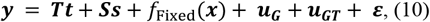

where ***u_GT_*** models the reaction norm for G×T effects as follows:

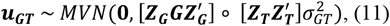

where ***Z_T_*** is the incidence matrix for the phenotype and trials. Thus, Eq. 11 allows for independent random effects between trials and has the potential to capture the unexpected G×T effects mentioned in the second assumption for genetic effects. Eqs. 8 and 10 can be combined as follows:

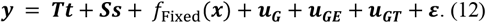

Eq. 12 can model both environment-specific and trial-specific random effects. The above random effect models are used in Jarquín *et al*. (2014) and Lopez-Cruz *et al*. (2015). The last random effect model is as follows:

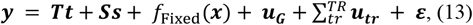

where *TR* is the total number of trials included in the phenotype data. 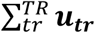 is represented as follows:

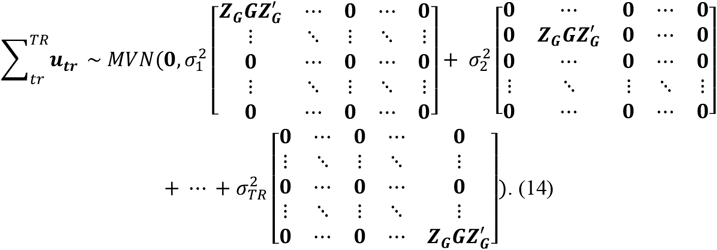

This random effect design is the same as that of Sousa *et al*. (2017) and Dahl *et al*. (2020), which allow for different heritability between trials. Therefore, the random effect model in Eq. 14 is more flexible and can capture G×T effects better than Eqs. 9 and 11. Information on the random effect terms is summarized in Table 1.

#### Fitting the LMMs

The LMMs described above include different numbers of random effect terms. For example, Eq. 7 includes only one random effect term (i.e., ***u_G_***) whereas Eq. 8 includes two (i.e., ***u_G_*** and ***u_GE_***). However, the model fitting in this study was performed using the same procedure. To explain the procedure, we generalized, tentatively, the random effects as follows:

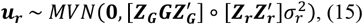

where ***u_r_*** is the *r*-th random effect, ***Z_r_*** is the incident matrix for phenotype and *r*-th random effects, and 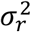 is the variance for ***u_r_***. To solve the LMMs with *R* random effects, we estimated the genetic variances without fixed QTL effect terms, as follows:

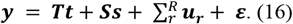

The solution of Eq. 16 was used to estimate a weight (*w_r_*) for each random effect term.

For example, the weight of the first random effect (*w*_1_) is calculated as follows:

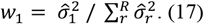

The variance–covariance matrix for the integrated random effects (***K***) was calculated as:

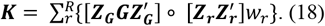

Next, the integrated random effects ***u_k_*** were derived as follows:

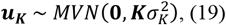

where 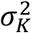 is the variance for ***u_K_***. Thus, the LMMs used to calculate the test statistics in the present study can be expressed as follows:

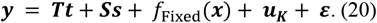

This method made GWAS computationally feasible for the present study. All calculations and parameter estimations for the LMMs were performed using the R package “gaston” (Perdry and Dandine-Roulland 2018).

#### Statistical tests for the fixed QTL effect terms

The statistical significance of the fixed QTL effect terms in each LMM (i.e., *f*_Fixed_(***x***)) was evaluated using the log-likelihood (LL) ratio test (LRT). The tests and formulae used to calculate the deviance and degrees of freedom are described in Table 2. The LRT used in this study can be divided into two categories (Table 2). The first category is a test for all QTL-effect terms (Table 2); it tests for the existence of a QTL in LD with the SNP. Thus, signals detected by this test can represent QTLs with or without Q×E effects. The second category is a test only for interaction terms (Table 2), which will be significant only when the QTL has Q×E effects (Malosetti *et al*. 2013). The LL of the maximum likelihood estimates was calculated using the *lmm.diago.profile.likelihood* function in the R package “gaston” (Perdry and Dandine-Roulland 2018). The *p* value of each test was calculated using the chi-square test based on the deviance and degrees of freedom (df) (Table 2).

**Table 2.**
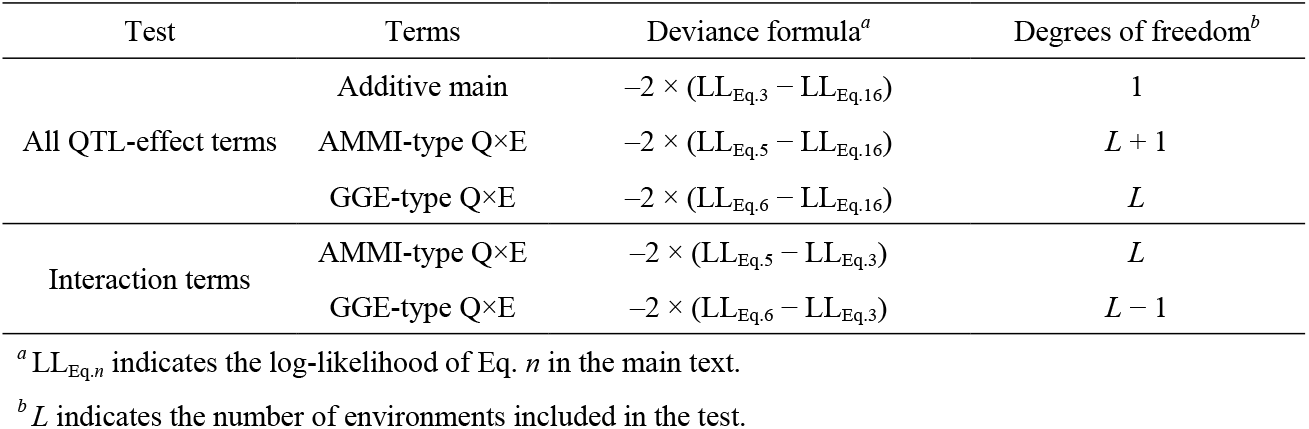
Hypotheses and terms of the chi-square tests performed in this study.

| Test | Terms | Deviance formula<sup>a</sup> | Degrees of freedom<sup>b</sup> |
| --- | --- | --- | --- |
| All QTL-effect terms | Additive main | $-2 \times (LL_{Eq.3} - LL_{Eq.16})$ | 1 |
| | AMMI-type Q×E | $-2 \times (LL_{Eq.5} - LL_{Eq.16})$ | $L + 1$ |
| | GGE-type Q×E | $-2 \times (LL_{Eq.6} - LL_{Eq.16})$ | $L$ |
| Interaction terms | AMMI-type Q×E | $-2 \times (LL_{Eq.5} - LL_{Eq.3})$ | $L$ |
| | GGE-type Q×E | $-2 \times (LL_{Eq.6} - LL_{Eq.3})$ | $L - 1$ |

<sup>a</sup> $LL_{Eq.n}$ indicates the log-likelihood of Eq. $n$ in the main text.
<sup>b</sup> $L$ indicates the number of environments included in the test.

Although the LRT can be used to obtain more accurate statistics in small- and moderate-sized samples, its drawback is the absence of test statistics for each coefficient included in the fixed effect terms that account for Q×E effects. As a complement to the LRT, we performed the Wald test for each coefficient for the Q×E effects. The Wald test scores were calculated as follows:

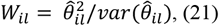

where 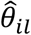 and *W_il_* indicate the estimated effect size of the *i*-th SNP in the *l*-th environment and the Wald score, respectively. Next, the *p* value of *W_il_* was calculated based on the chi-square test (df = 1).

We determined the genome-wide significant thresholds based on the false discovery rate (FDR), which is commonly applied in GWAS (Benjamini and Hochberg 1995; Storey and Tibshirani 2003). The FDR was calculated as described by Storey and Tibshirani (2003).

### Simulation experiments

We designed simulation experiments to evaluate the QTL detection power of the LMMs, under conditions described in detail in Table 3. The simulated major QTLs were randomly selected from the 16,782 SNP markers in a real dataset for tomatoes (Yamamoto *et al*. 2016). The simulated phenotypic values were generated using the following equation:

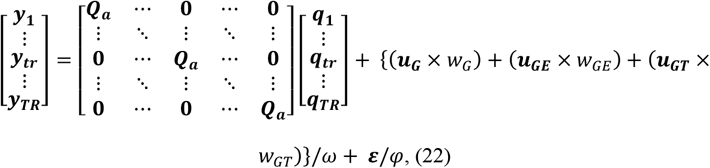

where ***y_tr_*** is a *m* × 1 vector for the phenotypic values in the *tr*-th simulated trial; ***Q_a_*** is an *m* × 3 matrix whose column elements consist of SNP genotypes selected for simulated QTLs; ***q_tr_*** is a 3 × 1 vector for the QTL effect in the *tr*-th trial. The elements of the ***q_tr_*** vectors become the same for trials under the same environmental conditions. (***u_G_*** × *w_G_*), (***u_GE_*** × *w_GE_*), and (***u_GT_*** × *w_GT_*) are random effects that follow Eqs. 4, 9, and 11, respectively. *ω* and *φ* were scalars required to adjust the phenotypic values to satisfy the given proportion of variance explained by each major QTL (PVE_QTL_) and heritability. In this study, the random effect values (i.e., ***u_G_, u_GE_***, and ***u_GT_***) were generated using the *mvrnorm* function in R using a corresponding variance–covariance matrix (R Core Team 2019). The relative contributions of the three random effects were adjusted by multiplying the values generated by *mvrnorm* and the specified weight parameters (*w_G_, w_GE_*, and *w_GT_* in Table 3). The residual values (***ε***) were generated using the *rnorm* function in R. In the present study, PVE_QTL_ and heritability for phenotypic values from all environments and trials were set to 0.1 and 0.5, respectively. To satisfy these settings, the optimal *ω* and *φ* were determined using the *optimize* function in R.

**Table 3.**
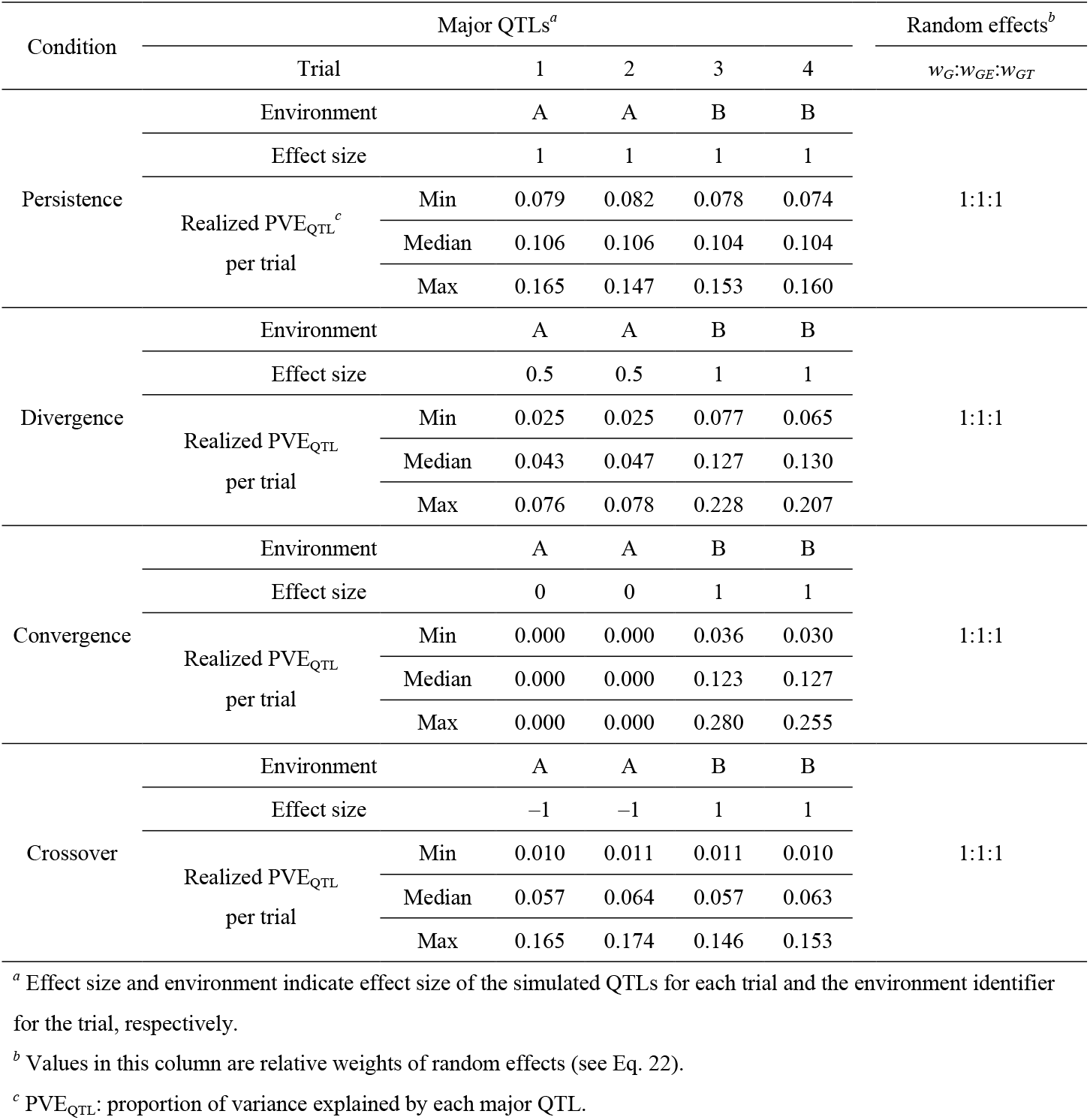
Parameters used for simulated phenotypes.

We assumed four Q×E effects: persistence, divergence, convergence, and crossover (Malosetti *et al*. 2013) (Table 3). Persistence means that the QTL has a persistent effect across environments and, therefore, has no Q×E effects. Divergence means that the QTL has different degrees of effect size between environments. Convergence means that the QTL shows an effect in a particular environment, but not in other environments. Crossover means that the direction of the QTL effect differs among environments.

### GWAS efficiency evaluation

We calculated recall, precision, and F-measure to evaluate GWAS power as in the previous studies (Hamazaki and Iwata 2020; Saber and Shapiro 2020; Shafquat *et al*. 2020). Recall is the proportion of true positives that are correctly identified; precision is the proportion of true positives among the retrieved positive signals; and F-measure represents the harmonic mean of recall and precision, calculated as follows:

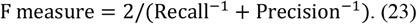

Thus, a high F-measure can only be achieved by balancing high precision and high recall. Generally, recall, precision, and F-measure are affected by the genome-wide significance threshold (Gage *et al*. 2018). Therefore, we also calculated the receiver operating characteristic (ROC) curve, and evaluated GWAS efficiency based on the area under the ROC curve (AUC) (Gage *et al*. 2018). An AUC of 0.5 indicates that the GWAS has no power to detect true signals, whereas an AUC of 1.0 indicates that all top signals of the GWAS agree with the true signals. We used the *roc* function in the “pROC” package (Robin *et al*. 2011) to calculate the AUC from −log_10_(*p*) values.

### Genomic inflation factor

Although the LMMs applied in GWAS are designed to avoid inflation of −log_10_(*p*) values due to population structure (Yu *et al*. 2008), the inclusion of too many covariates in an LMM often results in inflation or deflation of *p* values from the expected distribution (Moore *et al*. 2019). Therefore, the genomic inflation factor (*λ*_GC_; Devlin and Roeder 1999) is often used to evaluate the degree of *p* inflation or deflation (Voorman *et al*. 2011; Moore *et al*. 2019). In this study, *λ*_GC_ was calculated as follows:

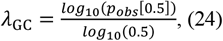

where *p_obs_*[0.5] indicates the 0.5 quantile of the observed *p* values. *λ*_GC_ > 1 indicates inflation of −log_10_(*p*) values and an increase in false positive signals, whereas *λ*_GC_ < 1 indicates deflation of −log_10_(*p*) values and an increase in the rate of false negatives (Devlin and Roeder 1999).

### Tomato genotype and phenotype data

We applied the LMMs to real tomato genotype and phenotype data (Yamamoto *et al*. 2016, 2017). We used 96 big-fruited tomato F_1_ varieties intended for the fresh market. These varieties were developed by various organizations such as seed companies and the public sector (Yamamoto *et al*. 2016). The parental combinations of the F_1_ varieties are unknown. Genotype data consisting of 16,782 SNP markers were obtained using Axiom myDesign genotyping arrays (Affymetrix Co., Ltd., Santa Clara, CA, USA). All SNP markers had a minor allele frequency of > 0.05, and a missing value rate of 0. Phenotyping was performed in the winter and spring cropping seasons in four and two trials, respectively. All plants were grown hydroponically using a high-wire system in a greenhouse at the National Agriculture and Food Research Organization at the Institute of Vegetable and Tea Science in Tsu, Japan. One plant per variety was grown in each trial. Among the phenotypes obtained (Yamamoto *et al*. 2016, 2017), we focused on the average fruit weight and fruit set ratio. The fruit set ratio indicates the ratio of flowers that reached fruit set. Values were transformed using the empirical logit transformation.

### Data availability

The phenotype and genotype data, R scripts, and R package developed for this study are available at https://github.com/yame-repos/gwasQxE.

## RESULTS

### Evaluation of power to detect Q×E effects

#### Common results among all simulation conditions

In LMMs including only the additive main effect term, there was little difference among the methods used to model random effects (Figure 2). The tests including Q×E effect terms showed different modes of action depending on the random effects. The random effects ‘***u_G_***’ and 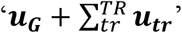 (Table 1) showed higher recall than other random effect models (Figure 2). Conversely, the precision, F-measure, and AUC of tests using ‘***u_G_***’ and 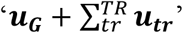 were lower than those of the other random effect models (Figure 2). These results indicate that signals detected in the tests that used ‘***u_G_***’ and 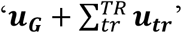 included more false discoveries than those in the other random effect models.

**Figure 2.**
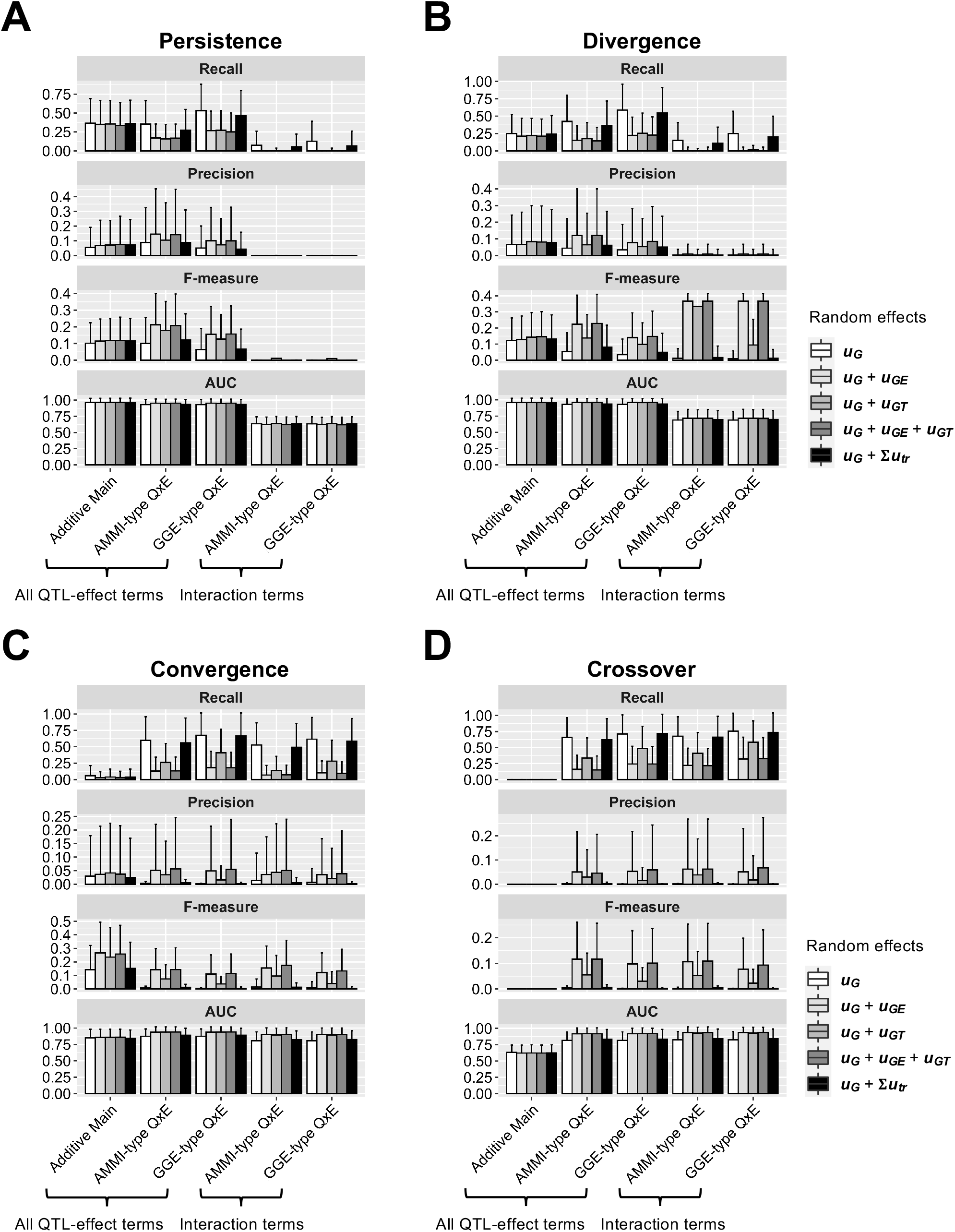
Bar plots of power to detect (A) persistence, (B) divergence, (C) convergence, and (D) crossover quantitative trait loci (QTLs) in the simulated phenotypes, under the assumption of multiple environments and multiple trials. Labels on the *x*-axes correspond to the tests described in Table 1 and 2. Recall, precision, and F-measure were calculated using a false discovery rate of 0.05 as the genome-wide significance threshold. Values represent means of 100 simulations.

#### Persistence

In this simulation, tests for interaction terms showed no power (i.e., recall ≈ 0 and AUC ≈ 0.5) (Figure 2A). This result is reasonable because no Q×E effects were included in the simulated QTLs (Table 3). LMMs including only the additive main effect were the fittest models for persistence QTLs. LMMs that included only the additive main effect showed the highest AUC, without obvious disadvantages in terms of recall, precision, and F-measure (Figure 2A). These results indicate that LMMs that include only the additive main effect are recommended for detecting persistence QTLs.

#### Divergence

The difference between the simulation conditions of ‘persistence’ and ‘divergence’ is that the latter QTLs have smaller PVE_QTL_ in environment A (Table 3). Because of the difference in realized PVE_QTL_ between environments, we expected LMMs that included Q×E effect terms to have higher power than those that included only the additive main effect. However, the results for ‘divergence’ resembled those for ‘persistence’ (Figure 2A and B). These results indicate that it is difficult to identify the ‘divergence’ effect using the LMMs in this study.

#### Convergence

Under this simulation condition, a QTL has an effect in environment B, but no effect in environment A (Table 3). Unlike persistence and divergence QTLs, LMMs that included only the additive main effect showed the lowest recall and AUC values (Figure 2C). This result indicates that Q×E terms are necessary to detect convergence QTLs. Another difference between ‘convergence’ compared to ‘persistence’ and ‘divergence’ is that the tests for interaction terms showed recognizable degrees of recall, precision, and F-measure (Figure 2C). These results indicate that the convergence Q×E effect can be detected using LMMs including Q×E fixed effect terms.

#### Crossover

Under this simulation condition, the QTL effect takes opposite directions in environments A and B (Table 3). LMMs with only an additive main effect term showed near-zero recall, precision, and F-measure (Figure 2D). These results indicate that Q×E terms are necessary to detect crossover QTLs. Recall, precision, F-measure, and AUC were equivalent between all QTL-effect terms and interaction terms (Figure 2D). These results indicate that crossover QTLs were detected as QTLs with Q×E effects and were not misrecognized as QTLs with only an additive main effect.

### Genomic inflation factor

The genomic inflation factor (*λ*_GC_) was calculated to assess the deviation of test statistics from the expected null distribution (Figure 3). Because we simulated only three major QTLs in the experiments, *λ*_GC_ should be equal to 1. Thus, the condition *λ*_GC_ ≈ 1 is necessary for precise calculation of FDR for a genome-wide significant threshold. In the tests for LMMs with only an additive main effect term (Table 2), – log_10_(*p*) values were close to the theoretically expected distribution (i.e., *λ*_GC_ ≈ 1) (Figure 3A). The tests for LMMs including Q×E effect terms yielded different distributions depending on the random effects. The *λ*_GC_ values were inflated and fluctuated when the random effects were ‘***u_G_***’ or 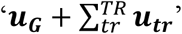 (Figure 3A). The use of the random effects ‘***u_G_ + u_GT_*** or ‘***u_G_ + u_GE_ + u_GT_***’ restrained *λ*_GC_ fluctuation, but the mean values differed depending on the fixed effect terms (Figure 3A). Figure 3B compares quantile–quantile (Q–Q) plots of mean −log_10_(*p*) values between LMMs with the random effects ‘***u_G_ + u_GE_ + u_GT_***’ (Table 2). The –log_10_(*p*) values of the AMMI-type Q×E effect terms showed deflation compared with the theoretically expected distribution (Figure 3B). These results indicate that GGE-type Q×E effect terms are more appropriate than the AMMI-type. Next, we focused on the relationship between GGE-type Q×E effect terms and the method used to model random effects. The random effects ‘***u_G_*’, ‘***u_G_ + u_GT_***’, and 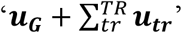 showed inflation of *λ*_GC_, whereas ‘***u_G_*** + *u_GE_*** and ‘***u_G_ + u_GE_ + u_GT_***’ showed *λ*_GC_ ≈ 1 (Figure 3C). These results indicate that an LMM that includes GGE-type Q×E effect terms and random effects ‘***u_G_ + u_GE_ + u_GT_***’ is recommended for calculating FDR as a genome-wide significant threshold.

**Figure 3.**
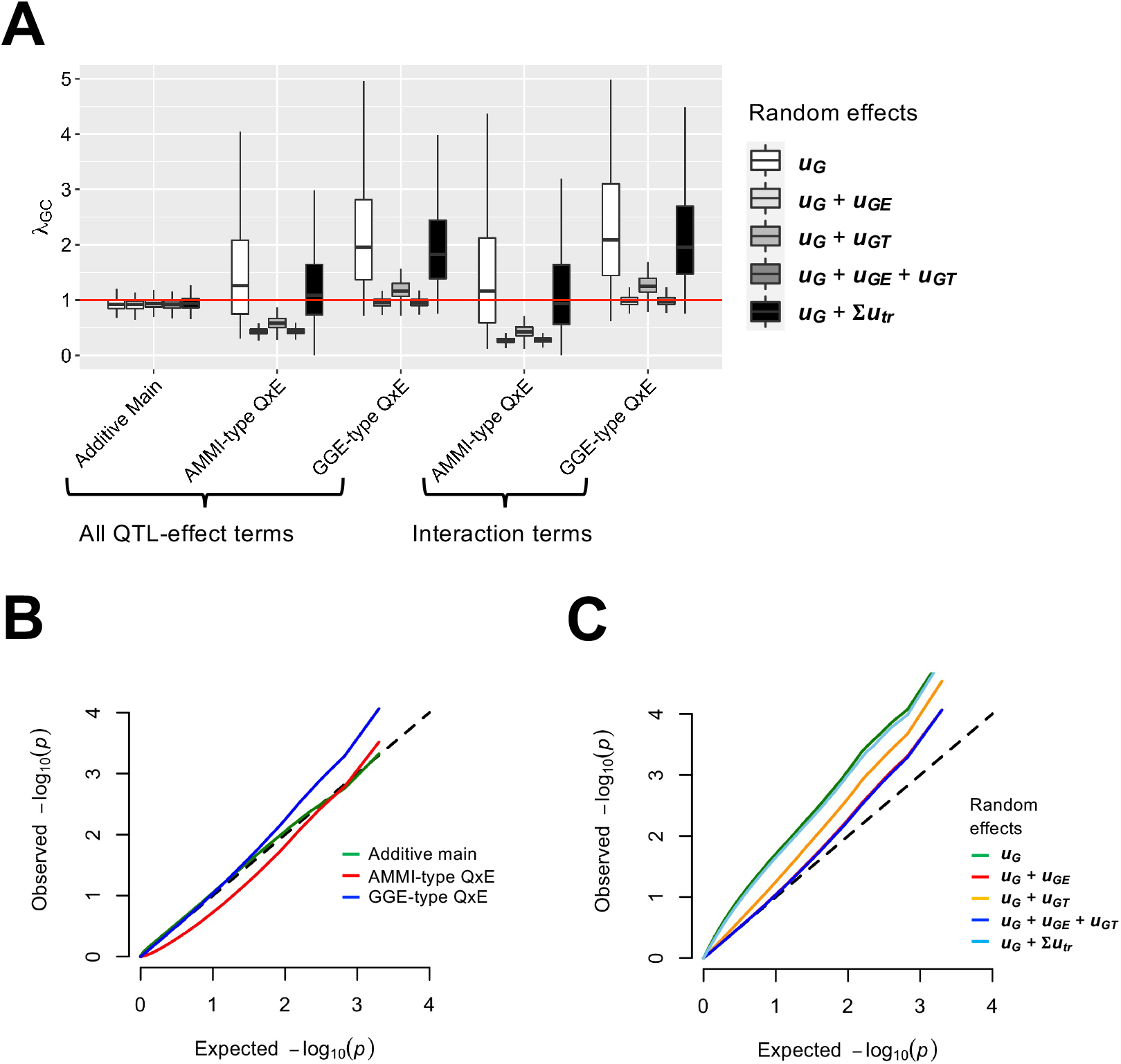
Evaluation of linear mixed models (LMMs) based on deviation of the *p* value distribution from the null hypothesis. (A) Box and whisker plots of genomic inflation factors for *p* values. Red horizontal line indicates the theoretically expected value (i.e., *λ*_GC_ = 1). (B) Quantile–quantile (Q–Q) plots of *p* values obtained from LMMs that include random effect ‘***u_G_ + u_GE_ + u_GT_***’. (C) Q–Q plots of *p* values obtained from LMMs with GGE-type Q×E fixed effect terms. In (B) and (C), *p* values shown in the panel represent means of 1,000 quantiles from 400 experiments (100 simulations × 4 conditions in Table 3), and a black dashed line indicates the *p* value distribution under the null hypothesis.

### GWAS for tomato agronomic traits

In the evaluation of the power to detect Q×E effects, LMMs that included only the additive main effect term were more efficient for persistence and divergence QTLs (Figure 2A and B). Conversely, LMMs including Q×E effects were necessary to detect convergence and crossover QTLs (Figure 2C and D). In the analysis using the genomic inflation factor, an LMM including GGE-type Q×E effect terms and the random effect ‘***u_G_ + u_GE_ + u_GT_***’ was recommended (Figure 3). Therefore, we used two LMMs for the tomato agronomic trait data. The first was an LMM that included only the additive main effect term and the random effect ‘***u_G_ + u_GE_ + u_GT_***’ (Table 1), and the second was an LMM that included GGE-type Q×E effect terms and the random effect ‘***u_G_ + u_GE_ + u_GT_***’ (Table 1). We focused on QTLs with FDR > 0.05, which are often used as genome-wide significant thresholds in GWAS (Cortes *et al*. 2020).

For average fruit weight, a significant signal on chromosome 9 was detected for the additive main effect and all QTL effect terms, including Q×E (Figure 4A and Table 4). The signal disappeared when the test was performed only for interaction terms (Figure 4A). These results indicate that the QTL detected on chromosome 9 is a persistence or divergence QTL. In the tests for interaction terms, we detected a significant signal on chromosome 10. The estimated effect size and the Wald score suggest that the signal is a convergence QTL that shows the effect only in the spring cropping season (Table 4).

**Figure 4.**
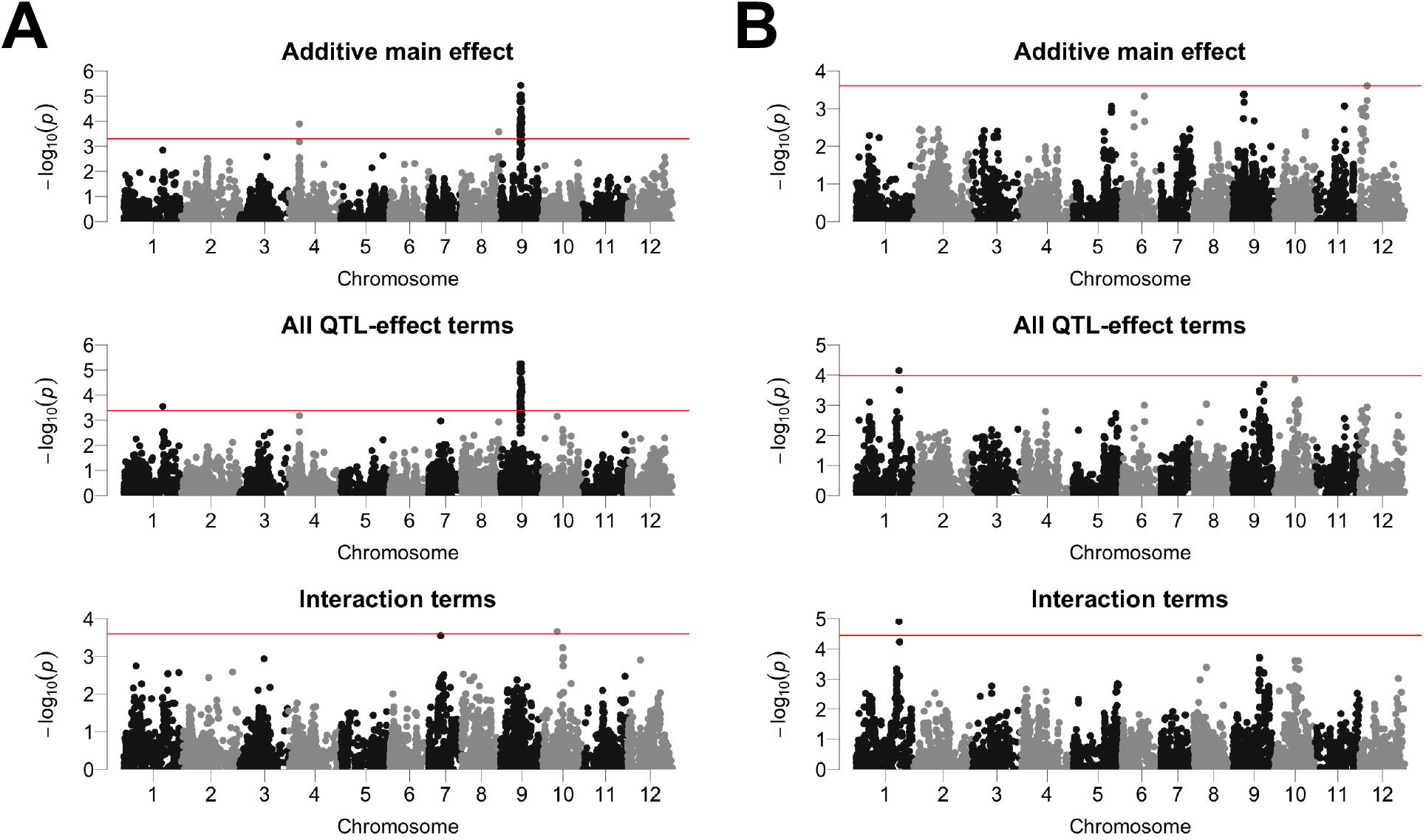
Genome-wide association study Manhattan plots for tomato agronomic traits. Red horizontal lines indicate the genome-wide significance threshold (false discovery rate < 0.05). (A) Average fruit weight. (B) Fruit set ratio.

**Table 4.**
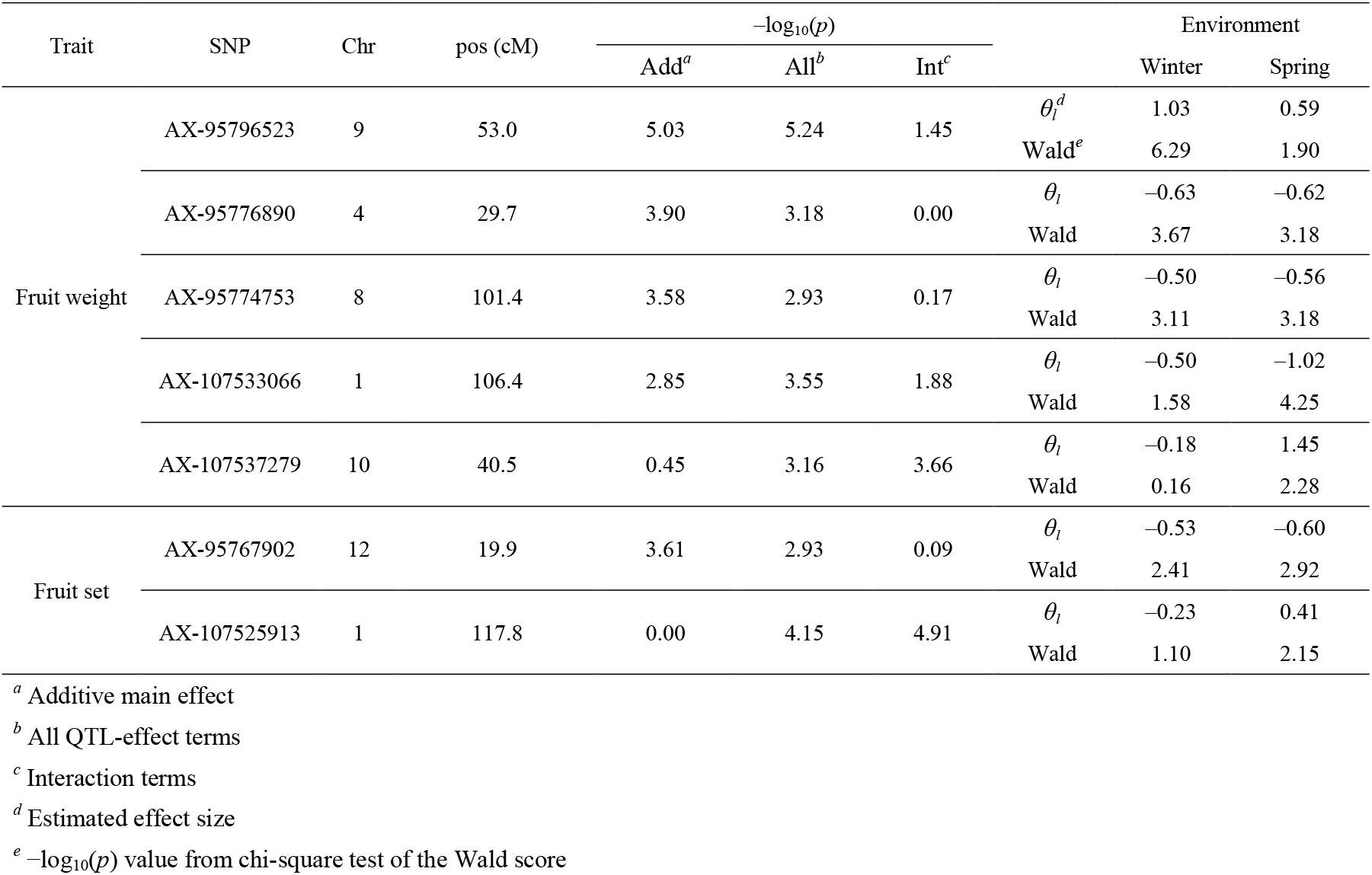
Statistics obtained in the genome-wide association study of tomato phenotypic data.

For fruit set ratio, a significant signal was detected on chromosome 12 in tests for the additive main effect (Figure 4B; Table 4). This signal disappeared in the other tests (Figure 4B). The effect size and Wald score of the signal on chromosome 12 were similar for the winter and spring cropping seasons (Table 4). These results indicate that the QTL on chromosome 12 is a persistence QTL. Alternatively, tests for all QTL effect terms, and tests for only interaction terms, detected significant signals on chromosome 1 (Figure 4B). The effect size of the signal was opposite in the winter and spring cropping seasons (Table 4). These results suggest that the QTL on chromosome 1 is a crossover QTL.

## DISCUSSION

In this study, we explored efficient LMMs to detect QTLs with Q×E effects. For this objective, we compared the efficacy of LMMs with various combinations of fixed QTL effect terms and random effect modeling methods (Tables 1 and 2). Efficacy was evaluated using recall, precision, F-measure, and AUC (Figure 2). None of the tested LMMs showed high values for all parameters (Figure 2). Notably, LMMs with higher recall showed lower precision and F-measure (Figure 2). This result is reasonable because recall and false positives are highly correlated in GWAS (Bian and Holland 2017). Generally, recall, precision, and F-measure change depending on the level of the genome-wide significance threshold (Gage *et al*. 2018). These problems make it difficult to select the most efficient LMM. Recently, AUC has been used to evaluate GWAS efficiency because it is independent of the genome-wide significance threshold (Gage *et al*. 2018; Moore *et al*. 2019; Hamazaki and Iwata 2020; Shafquat *et al*. 2020). In the present study, AUC values were affected by random effect modeling methods to a smaller degree than recall, precision, and F-measure (Figure 2). Therefore, the AUC was useful for selecting the most effective LLM in this study.

Inflation or deflation of *λ*_GC_ causes various problems in GWAS (Devlin and Roeder 1999). For example, although FDR is commonly applied to determine genome-wide significant thresholds in GWAS (Benjamini and Hochberg 1995; Storey and Tibshirani 2003), its calculation requires a theoretically expected *p* value distribution (i.e., *λ*_GC_ ≈ 1). Genomic control (GC) corrects inflated or deflated *p* values using the *λ*_GC_ value (Devlin and Roeder 1999). However, the applicability of a given inflation factor (i.e., *λ*_GC_) to correct marker *p* values differs according to allele frequency and correlation with other covariates, and therefore use of a uniform overall inflation factor (i.e., *λ*_GC_) may results in a loss of power (Price *et al*. 2006; Wang *et al*. 2012; Moore *et al*. 2019). Notably, *λ*_GC_ ≈ 1 is theoretically correct for the present study because we simulated only three major QTLs (Table 3) and, therefore, most *p* values should follow the expected distribution. In this study, the use of AMMI-type Q×E effect terms resulted in deflated *λ*_GC_ (Figure 3), although the AUC values were equivalent to those of the GGE-type Q×E effect (Figure 2). This result is attributed to the df of the LRT (Table 2). Because of the additive main effect term (***x**β* in Eq. 5), the LRT that used AMMI-type Q×E effect terms had one more df than that using GGE-type Q×E effect terms (Table 3). Therefore, we concluded that using GGE-type Q×E effect terms is preferable to GWAS for Q×E effects. However, a desirable feature of the AMMI-type Q×E effect terms is the separation of a QTL effect into additive main and Q×E effects. To perform exhaustive Q×E effect analysis with minimal oversight, our findings suggest that (1) genome-wide analysis with LMMs should be performed using GGE-type Q×E effect terms, and (2) Q×E effects should be assessed using AMMI-type Q×E effect terms only for significant signals detected in (1).

In this study, the random effect 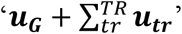 showed low precision, F-measure, and AUC values (Figure 2) and was less effective in controlling *λ*_GC_ (Figure 3), although its efficacy has been demonstrated in a previous study (Sousa *et al*. 2017). One possible reason for this discrepancy is that of differences in methods used to estimate the variance component in LMM fitting. In this study, we performed direct estimation using the average information restricted maximum likelihood method without fixed QTL effect terms (Gilmour *et al*. 1995; Perdry and Dandine-Roulland 2018). Conversely the variance component estimation and fitting method used in Sousa *et al*. (2017) were based on a Bayesian approach that updates the parameters using a Gibbs sampler (Pérez and de Los Campos 2014). Because Sousa *et al*. (2017) performed model fitting to construct prediction models for GS, one model fitting process per trait was sufficient. Conversely, GWAS must perform model fitting for every marker in the data, which can be computationally unfeasible under the Bayesian approach. Therefore, we did not use the Bayesian approach for model fitting in this study, even though it has the potential to increase the power of QTL detection.

The findings of the present study can be summarized as follows. First, LMMs including Q×E effect terms are necessary to detect QTLs with Q×E effects (Figure 2). Second, the random effect ‘***u_G_ + u_GE_ + u_GT_***’ is necessary to control *λ*_GC_ and calculate FDR appropriately (Figure 3). We applied these findings to real phenotypic data for tomatoes to detect QTLs associated with agronomic traits. A fruit weight QTL on chromosome 9 was located on the *fw9.1* region, which was identified in a genetic mapping population derived from a cross between cultivated and wild tomato (Tanksley *et al*. 1996). A fruit set QTL on chromosome 1 was close to *fset1.3*, which was detected in a genetic mapping population derived from eight tomato varieties (Diouf *et al*. 2020). Interestingly, Diouf *et al*. (2020) suggested that*fset1.3* has Q×E effects, which is consistent with the results of this study (Figure 4 and Table 4). Thus, the findings of this study have the potential to contribute to the identification of more QTLs with Q×E effects.

## ACKNOWLEDGMENTS

This work was supported by the Japan Science and Technology Agency (JST) PRESTO (Grant number, JPMJPR16Q9), JSPS KAKENHI (Grant number, 19H02951), and Meiji University (Interdisciplinary Research Program in Mathematical Modeling and Life Sciences) to E.Y.

## APPENDIX

### Dominant effect QTLs

Because the materials used in this study were F_1_ varieties, we extended the analyses in the main text to include dominant effect QTLs. In this appendix, we briefly summarize the methods and results of this analysis.

#### LMMs

To include dominant effects, Eq. 1 was extended as follows:

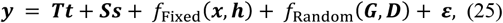

where *f*_Fixed_(***x, h***) and *f*_Random_(***G, D***) are the fixed and the random effect terms that include dominant effects, respectively. ***h*** is an *n* × 1 vector of SNP genotype heterozygosity coded as {0, 1, 0} = {aa, Aa, AA}. ***D*** is the dominance covariance matrix between individuals *j* and *k*(*D_jk_*), and was defined as follows:

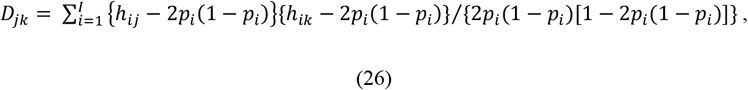

where *h_ij_* is the coded SNP genotype heterozygosity value for the *i*-th SNP of the *j*-th individual. The derivation of ***D*** is explained in detail in Su *et al*. (2012). Without consideration of differences among environments, the random effects for ***D*** are modeled as follows:

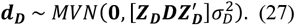

We omitted the derivation of dominant random effects for G×E (i.e., ***d_D_, d_DE_, d_DT_***, and 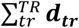) because these are analogous to the equations used for additive G×E random effects (i.e., ***u_G_, u_GE_, u_GT_***, and 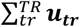). When the dominant effect terms were included, Eq. 3 was extended as follows:

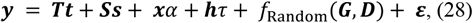

where *τ* is the dominant effect for a QTL in LD with the SNP. An LMM including the AMMI-type Q×E effect terms (Eq. 5) was extended as follows:

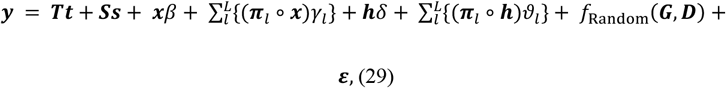

where *δ* is the dominant QTL effect not specific to the environment, and *ϑ_l_* is the dominant QTL effect specific to the *l*-th environment. An LMM including the GGE-type Q×E effect terms (Eq. 6) was extended as follows:

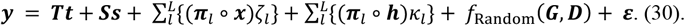

where *κ_l_* is the dominant QTL effect in the *l*-th environment.

#### Simulation conditions

The equation used to simulate phenotypic values, including dominant effects, was as follows:

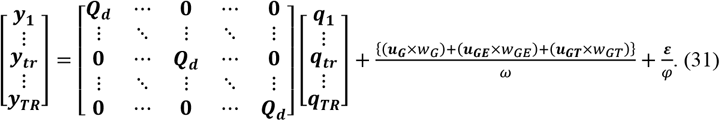

The difference between Eqs. 22 and 31 is that the coding of the SNP genotype values ***Q_a_*** in Eq. 22 is {0, 1, 2} = {aa, Aa, AA}, whereas that of ***Q_d_*** in Eq. 31 is {0, 3, 2} = {aa, Aa, AA}. This dominant effect design was based on a real dominant-effect QTL in tomatoes (Krieger *et al*. 2010).

#### Results and Discussion

The results obtained using the genomic inflation factor (*λ*_GC_) were similar to those of LMMs without dominant effects, except that the values were slightly deflated in all tests (Figure S1A), which was likely to be because the additional fixed terms in the dominant effect LMMs resulted in larger df in the subsequent chi-square test. The inclusion of dominant effect terms in the LMMs showed an advantage in terms of both recall and AUC for detecting the dominant-effect QTLs simulated in this study (Figure S1B). This result is reasonable because models including dominant effect terms generally provide a more effective fit to phenotypic values with a dominant effect (Letter *et al*. 2007). Nevertheless, LMMs that included dominant effect terms did not detect dominant-effect QTLs for the real agronomic trait data used in this study (Figure S2). One possible reason for this result is that the deflation of −log_10_(*p*) values, as indicated in *λ*_GC_ (Figure S1A), resulted in false negative QTLs. Another possibility is that, as suggested in some previous studies, the contribution of dominant effects is less evident for quantitative traits, and the use of an additive effect model is sufficient for genome-wide analysis (Varona *et al*. 2018). In addition, another problem in the inclusion of dominant effect terms have been discussed by Balding (2006). That is, the existence of few homozygotes for an allele results in the neglect of the homozygote genotypes and linear regression fitting between homozygote for the another allele and heterozygote genotypes (Balding 2006). Because the materials used in the present study were F1 varieties that include many genetic loci where heterozygous genotypes are major and homozygous are minor, this scenario is possible.

**Figure S1.**
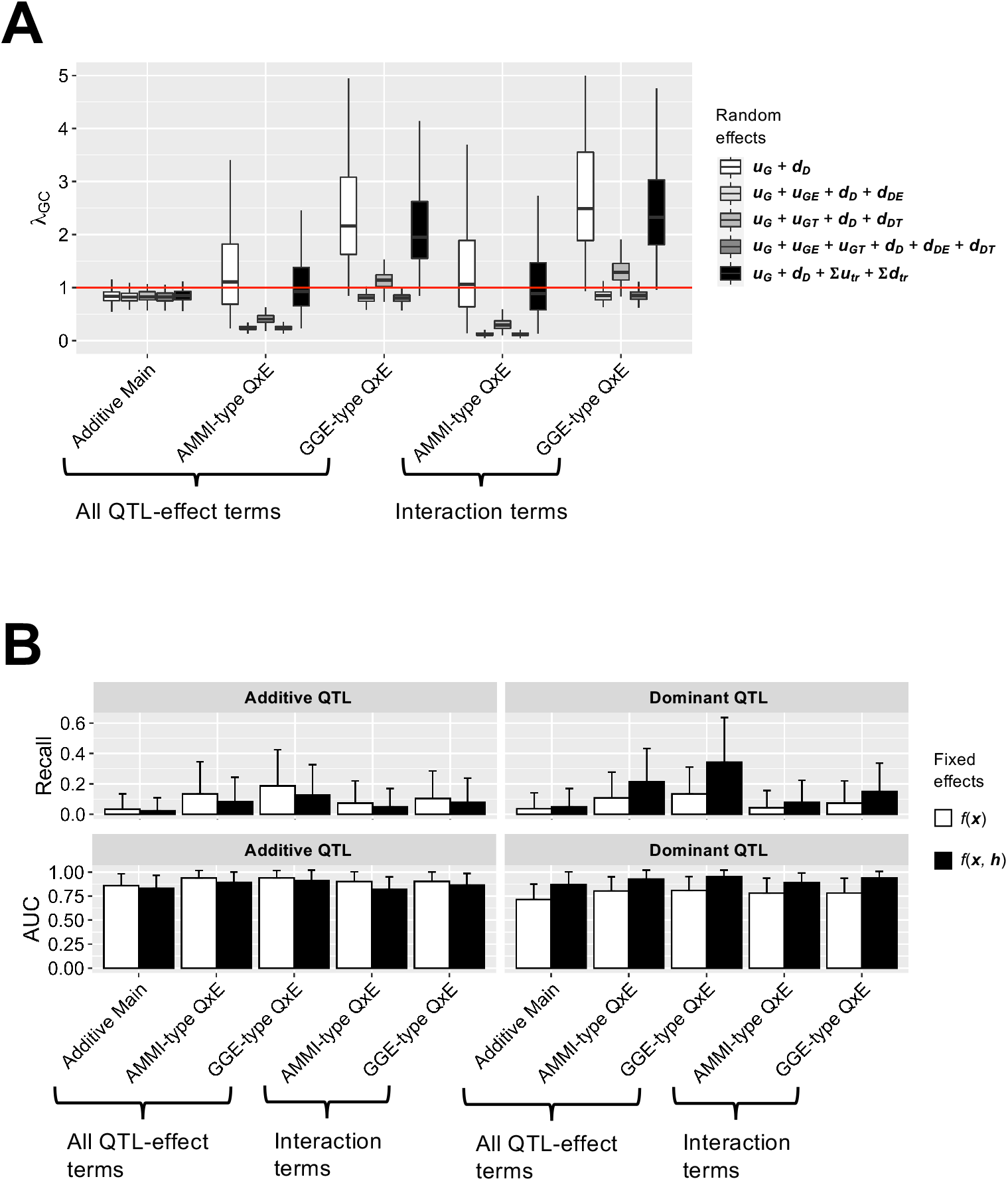
Evaluation of linear mixed models (LMMs) including dominant effect terms. (A) Box and whisker plots of genomic inflation factors (*λ*_G_C) for *p* values. Red horizontal line indicates the theoretically expected value (i.e., *λ*_GC_ = 1). (B) Bar plots of power to detect quantitative trait loci (QTLs) in simulated phenotypes, assuming multiple environments and multiple trials. Recall values were calculated using a false discovery rate of 0.05 as the genome-wide significance threshold. Values represent means of 100 simulations. The fixed and random effect terms used in this analysis was GGE-type Q×E and ‘***u_G_ + u_GE_ + u_GT_ + d_D_ + d_DE_ + d_DT_***,’ respectively.

**Figure S2.**
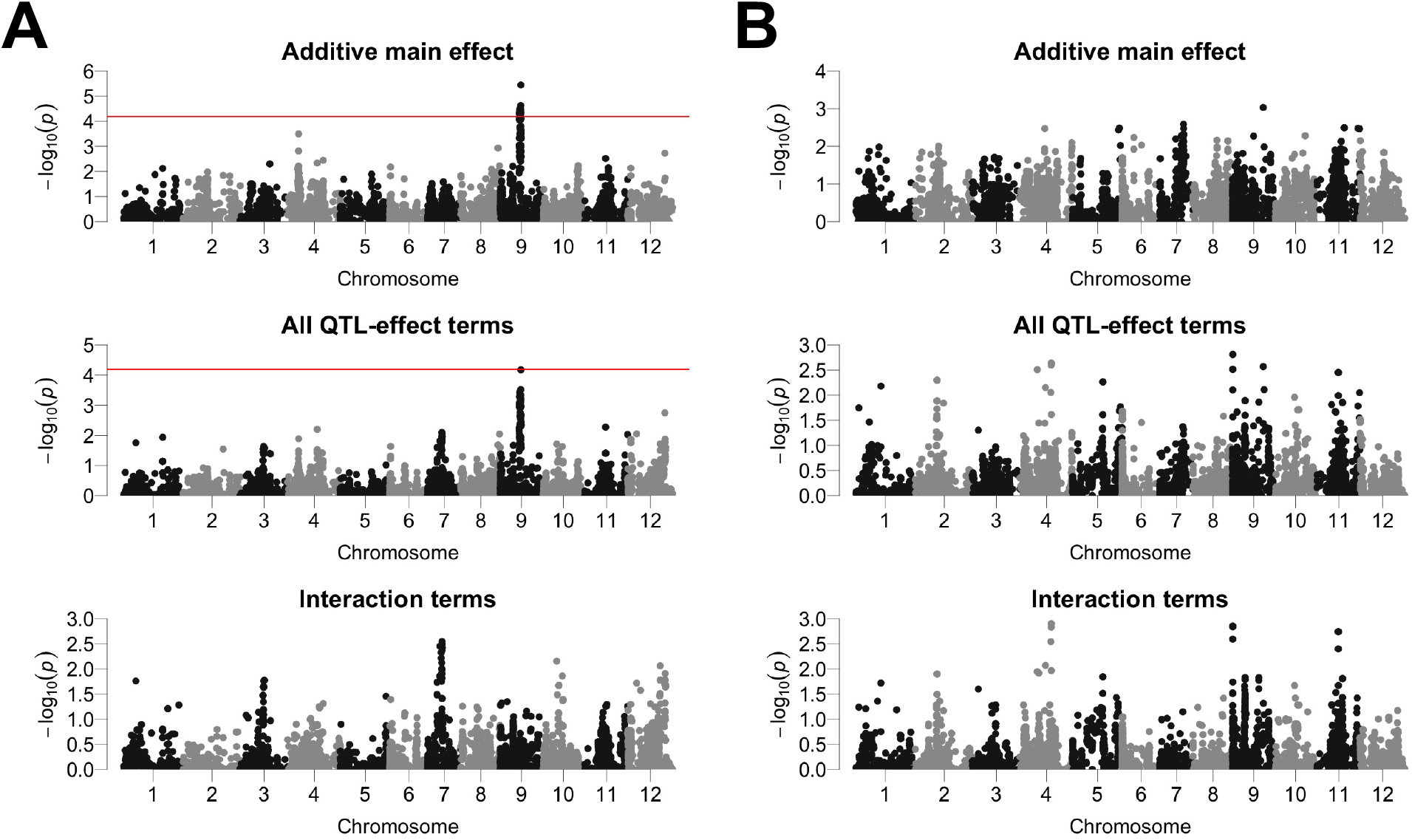
Genome-wide association study Manhattan plots of tomato agronomic traits. Red horizontal lines indicate the genome-wide significance threshold (false discovery rate < 0.05). (A) Average fruit weight. (B) Fruit set ratio.

